# Collagen-based bilayered biomimetic tubular materials for vascular and airway applications

**DOI:** 10.64898/2026.03.20.713181

**Authors:** Florian Fage, Alshaba Kakar, Ilaria Onorati, Isabelle Martinier, Alessia Castagnino, Dorian Verscheure, Emeline Saindoy, Othmane Darouich, Julien Gaudric, Valérie Besnard, Abdul I. Barakat, Emmanuel Martinod, Carole Planes, Nicolas Dard, Francisco M. Fernandes, Lea Trichet

## Abstract

Biomimetic tubular scaffolds hold great promise for tackling unmet clinical needs thanks to their biocompatibility and recapitulation of cellular microenvironments, conferring the ability to promote regeneration. Potential applications include small-diameter vascular implants and grafts for airway repair, for which no viable off-the-shelf solutions currently exist. The tubular materials (4 and 8 mm internal and external diameters) presented here consist purely of type I collagen, contain no chemical crosslinkers, and reproduce the multi-scale architecture of the native tissue including the presence of collagen fibrils. A novel two-step protocol provides materials with distinct concentric layers. A porous external structure, obtained by means of ice templating combined with collagen topotactic fibrillogenesis, favours oriented cell colonization. A smooth and much less porous internal layer provides mechanical and water-tightness properties relevant for *in vivo* implantation and promotes the formation of an endothelial monolayer under both static and flow conditions. The compliance of the double-layered materials under physiological pressure is close to that of piglet carotid arteries. The materials are also determined to be sufficiently flexible to provide the ability to perform *ex vivo* anastomosis with bronchi, although the relatively low value of suture retention strength remains a limitation for *in vivo* suturing.

## 1. Introduction

The design and implementation of biomaterials intended to replace tubular tissues such as small-diameter blood vessels and airways remain an unaddressed clinical challenge. These biomaterials need to exhibit several specific features including 3D organization in distinct concentric layers, water-tightness, and physiologically relevant mechanical properties. In these constructs, precise modulation of parameters such as composition, structure and surface topography is critical for enhancing cell adhesion, proliferation and differentiation.^[1]^ For instance, in the case of blood vessels, an ideal replacement would replicate the three-layer organization of the vascular wall: the tunica intima, consisting of a monolayer of endothelial cells anchored to a basement membrane, the tunica media, consisting in a fibrous extracellular matrix (ECM) containing collagen, elastin and proteoglycans with embedded smooth muscle cells (SMCs), and the external layer, or tunica adventitia.

Small-diameter (<6 mm) vessel pathologies include coronary heart disease (CHD) and peripheral arterial disease (PAD). In the most severe cases, revascularization techniques rely on vascular bypass grafting, with use of either autologous or synthetic materials.^[2]^ However, use of saphenous vein or internal thoracic artery grafts might not be possible due to comorbidities, while use of non-biodegradable polymer materials in small diameter vessels often results in thrombotic occlusion, manifesting itself usually within 5 years following surgery. In the case of airway transplantation, synthetic prosthetics often have some limitations, such as inflammatory granuloma obstructing the lumen, infection, and graft migration.^[3–5]^ Tracheal and bronchial allografts have limited availability, can lead to size mismatch, and require use of immunosuppressive drugs. Cryopreservation, lyophilization and fixation can spare the use of immunosuppressive treatment, but insufficient graft revascularization remains a major problem. Recently, the group led by Prof. Martinod demonstrated that airway bioengineering using cryopreserved and stented aortic allografts is a feasible approach for complex tracheal and bronchial reconstruction.^[6–8]^ *De novo* generation of cartilage was observed after a few months, reconstituting an alternate structure of cartilage rings, as well as regeneration of a mixed respiratory neoepithelium-like structure. Still, lack of donor tissue prevents this strategy from wider dissemination, and there remains an acute need for off-the-shelf biomimetic materials.

Type I collagen is the most abundant protein present in the ECM of mammals. In its tightly regulated fibrillated form, it provides structural support for cells, exhibits bioactive properties and, together with embedded cells and other components of the ECM, regulates the mechanical performance of the tissues. Type I collagen also has very low immunogenicity and is therefore a material of choice for the development of scaffold-based grafts.^[2]^ In these materials, recapitulation of tissue-specific native architectures and fibrillar structures is thought to be key for obtaining adequate mechanical properties and proper cellular colonization.^[9]^

Synthesis of collagen-based tubular materials relies on diverse techniques. Weinberg and Bell pioneered the gel-based approach to develop vascular models by casting a suspension of cells in a collagen solution inside a tubular mold, the resulting gel being further compacted though a cell-mediated process.^[10]^ This technique has demonstrated striking results in preclinical studies, with functional restoration and vascular tissue recapitulation with the three different layers;^[11]^ however, mechanical reinforcement by synthetic sleeves is required. Combining collagen with fibrin subsequently enabled the tuning of gel compaction and mechanical properties.^[12]^ Assembling of collagen and SMCs in a 3D cylindrical geometry followed by culture in a bioreactor enabled reaching higher mechanical properties and the combination of up to three layers of materials.^[13,14]^

Cell-free strategies have been developed in order to shorten fabrication time and to enhance collagen concentration and mechanical properties. Dehydration followed by crosslinking was introduced to obtain dense tubular constructs with enhanced mechanical properties.^[15]^ Dense collagen gel sheets obtained after plastic compression and composite matrices combining dried collagen gels and elastin were rolled around a cylindrical mandrel.^[16,17]^ In the latter case, the obtained vascular grafts demonstrated mechanical properties close to those of native tissues, with encouraging results obtained *in vivo*.^[17]^ Rotation-based water removal processes have enabled the development of tubular collagen gel scaffolds with moderate elastic moduli.^[18]^ This technique has also allowed obtaining trilayered arterial wall models with precise composition of embedded cells and endothelialization, and further maturation for 2 weeks led to improved mechanical properties along with compaction of tube walls.^[19]^ Justin *et al.* developed a method to densify collagen hydrogels while maintaining the thickness of the tubular construct walls, by withdrawing water from the gel *via* hydrophilic nylon membranes.^[20]^ This densification process is also cell-compatible and enables obtaining tubes with distinct cell populations in specific locations.

Despite these advances, current techniques do not allow coupling of collagen fibrillogenesis with internal structuration. In particular, macroporosity is an essential feature needed to ensure cellular homeostasis, by allowing diffusion of nutrients and waste and promoting cell guidance for proper host cell colonization and vascularization inside the material. Self-standing porous collagen materials can be obtained by freezing and lyophilization of a collagen solution with subsequent physical or chemical cross-linking.^[21,22]^ The technique was adapted to provide tubular porous materials displaying three layers.^[23]^ Thrombus formation observed one month after implantation was related to the porosity of the inside wall. Crosslinked atelocollagen solutions were also frozen and lyophilized to fabricate ring-shaped scaffolds alternately stacked with mimetics of native cartilaginous rings, namely chondroitin-sulfate-incorporating type-II atelocollagen scaffolds seeded with bone marrow stem cells.^[24]^ After submuscular implantation for 4 weeks in autologous rabbits, this biomimetic trachea demonstrated ample vascularization of the collagen I scaffold regions, and after tracheal implantation better results in terms of survival, airway patency and epithelialization were obtained when compared to tubes made of cartilaginous rings alone. Khalid *et al.* combined a 3D printed PCL framework with the previously developed bilayered collagen-hyaluronic acid scaffold, comprising a film and an interconnected porous sublayer obtained by freeze-drying, in order to reinforce mechanical properties.^[25,26]^ The process produced scaffolds compatible with growth of human umbilical vein endothelial cells (HUVECs) and mesenchymal stem cells (hMSCs) and that could be vascularized *ex vivo* in a chick chorioallantoic membrane model. In a subsequent study, spatial control of cell seeding was achieved, with the inner layer seeded using respiratory epithelial cells to mimic the tracheobronchial respiratory epithelium, while the outer layer was seeded using lung-derived fibroblasts.^[27]^

Collagen-based porous materials can also be obtained by means of biological textile approaches, which was first pioneered by Cavallaro *et al.*, with the use of extruded, dehydrated and cross-linked collagen threads.^[28]^ Electrochemically aligned collagen (ELAC) fibers were also used to knit a tubular graft.^[29]^ While the mechanical properties were highly satisfactory, the elevated pore size required electrospinning of collagen nanofibers in the luminal surface. Magnan *et al.* also showed that the textile approach might provide human watertight textiles with high mechanical strength, while still requiring a cell-mediated production step of 6 to 12 weeks, followed by yarn fabrication.^[30]^

Some of these latter observations highlight the importance during materials fabrication of both the collagen assembly process and precise control over graft topography for successful graft implantation. Indeed, a smooth surface of limited porosity is required for proper cellular lining on the internal wall and recapitulation of a functional endothelium or epithelium, while an oriented porous structure in the external part of the material would allow rapid cell colonization from the outside.^[1]^

Freeze casting, a technique initially developed for the elaboration of porous ceramics and adapted to produce scaffolds based on biopolymers, offers the possibility to precisely tune material porosity by adjusting different parameters such as solution composition, freezing kinetics, and the temperature gradient.^[31–33]^ Recently introduced coupling of freeze casting with topotactic fibrillogenesis enables self-assembly of type I collagen into fibrillar constructs, while simultaneously stabilizing the macroscopic patterns defined by the ice templating technique. Precisely shaped porous materials are thus obtained in non-denaturing conditions.^[34,35]^ The obtained local concentration of collagen inside the pore walls is greatly enhanced and is in the same range as that observed in native tissues. The lyotropic nature of collagen thus gives rise to highly organized domains, and, upon fibrillogenesis, the multiscale hierarchical architecture of collagen is recapitulated, which provides materials with enhanced mechanical properties that are also prone to cell colonization. Ice templating coupled to topotactic fibrillogenesis also provides tubular materials whose macroscopic porosity can be modulated by the thermal conductivity of cylindrical molds, resulting in a new family of materials with tunable textural features and supramolecular arrangement of type I collagen.^[36]^

In this work, we propose to combine the obtained tubular external porous layer with an inner non-porous layer in order to achieve collagen-only biomimetic tubular materials that exhibit water-tightness and mechanical properties compatible with surgical manipulation. Colonization by endothelial cells and human mesenchymal stem cells was also investigated in the different layers to assess the potential for clinical applications.

## 2. Results and Discussion

### 2.1. Fabrication and structural characterization of sequentially assembled bilayered tubular collagen materials

Collagen-only tubular scaffolds composed of a porous outer layer and a compact luminal layer forming a smooth, dense internal surface were developed. The tubular constructs were fabricated through a two-step sequential process, consisting of the formation of a porous external tube with subsequent addition of a dense internal layer. These two layers were seamlessly integrated into a single cohesive material, resulting in tubular scaffolds exhibiting spatial variations in structure and collagen organization across the wall thickness. A schematic overview of the sequential fabrication process is provided in Figure 1.

**Figure 1.**
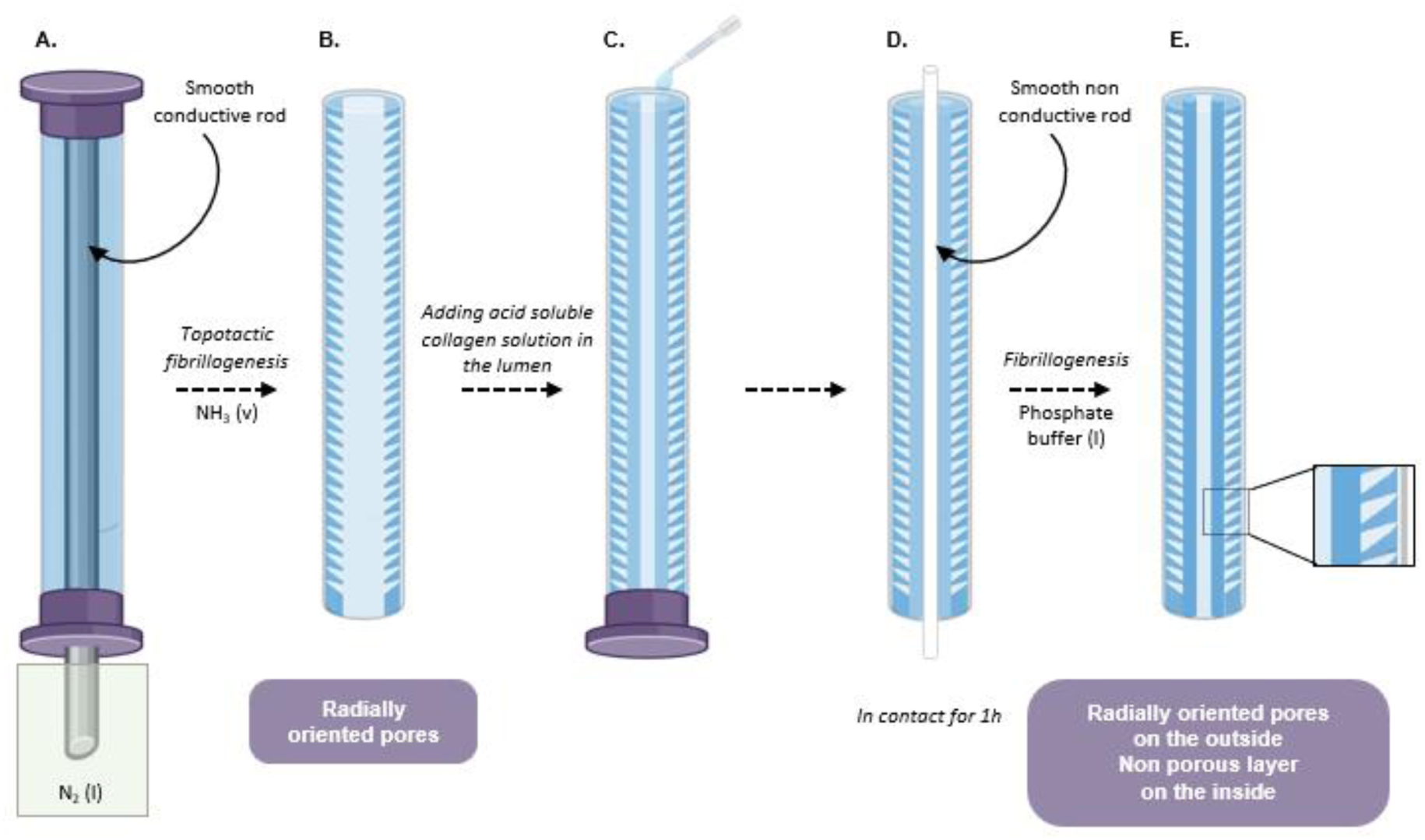
Schematic depicting the experimental design with the different steps leading to bilayered collagen-based tubular materials.

Briefly, controlled fabrication of the tubular geometry relied on a modular coaxial cylindrical mold system designed to independently define the external diameter, lumen size, and the respective thicknesses of the two layers. This process enabled fabrication of bilayered collagen tubes of 4- and 8-mm internal and external diameters, with control of the thicknesses of the porous outer and dense luminal layers of respectively 1.5 and 0.5 mm (Figure 2 A-B).

In practice, the porous outer tube was first fabricated using an external cylindrical shell defining the outer diameter of the tube used in combination with an initial internal rod. Structuration of this porous outer layer relied on freeze casting and consisted of plunging a type I collagen solution (40 mg/mL), placed between adequate tubular molds, inside liquid nitrogen at a controlled velocity (Figure 1A).^[36]^ In the present radial mold configuration, the external non-conductive PTFE tube and the internal conductive aluminum tube impose a radial thermal gradient, causing freezing to proceed preferentially from the inner conductive boundary toward the outer region, with radial growth of ice crystals across the tube wall.^[36,37]^ This process encoded a network of radially oriented, tunnel-like channels, as evidenced by scanning electron microscopy images obtained after lyophilization of the materials, directly reflecting the direction of ice crystal growth during freezing (Supplementary Figure 1). Subsequently, to stabilize the freeze-casted scaffold without altering its architecture, collagen fibrillogenesis was induced using cold ammonia vapor exposure followed by rinsing with water vapor, following the topotactic fibrillogenesis protocol (Figure 1B).^[34]^ The radially oriented tunnel architecture generated during freeze casting was thereby preserved after fibril formation, yielding a mechanically stable yet highly organized porous collagen scaffold. Once this porous tube was formed and stabilized, the initial internal rod was removed and replaced by a second internal rod of smaller diameter, and a dense luminal layer was formed by depositing a concentrated acidic collagen solution (26 mg/mL) onto the inner surface of the fibrillated porous scaffold (Figure 1C). The collagen solution was thus geometrically confined between the inner surface of the porous scaffold and the second internal tube, and the assembly was left to rest for 1 h, enabling the formation of a cohesive bilayered structure (Figure 1D).

The acidic collagen in contact with the fibrillated porous collagen layer induces local reorganization and partial redissolution of the superficial fibrillar network at the interface. During the resting period, this process allows molecular interpenetration between the newly deposited collagen molecules and the underlying fibrillated scaffold. Subsequent fibrillogenesis under high ionic strength solution (5X PBS) stabilized this reorganized region, resulting in a continuous and mechanically cohesive junction between the dense luminal layer and the porous outer layer (Figure 1E). This mechanism led to the formation of a unified collagen tube rather than a simple stacked assembly of two independent layers (Figure 2C-D). Confocal microscopy images show porosity in the outer layer in contrast to the smooth, compact, and non-porous inner surface. Porosity quantification on transversal section enables to determine a median width of the pores and of the pore walls of 14 ± 6 µm and 15 ± 6 µm, respectively, with a porous fraction of 52% (data not shown). Both layers are fused all along the longitudinal (Figure 2C) and transversal (Figure 2D) sections. The resulting scaffold thus combines a highly organized radial architecture with a mechanically stable fibrillar collagen matrix. Transmission electron microscopy examination confirmed the presence of a fibrillar collagen network throughout the tube wall. Tightly packed and aligned small fibrils (Figure 2E-F), exhibiting a wave or arch pattern, correspond to the preservation of the liquid-crystal organization fibrillogenesis, with cholesteric phase, as already observed inside the walls of the porous materials.^[34,35,38]^ In the non-porous region, randomly oriented collagen fibrils present the characteristic 67 nm D banding (Figure 2G).^[35]^

**Figure 2.**
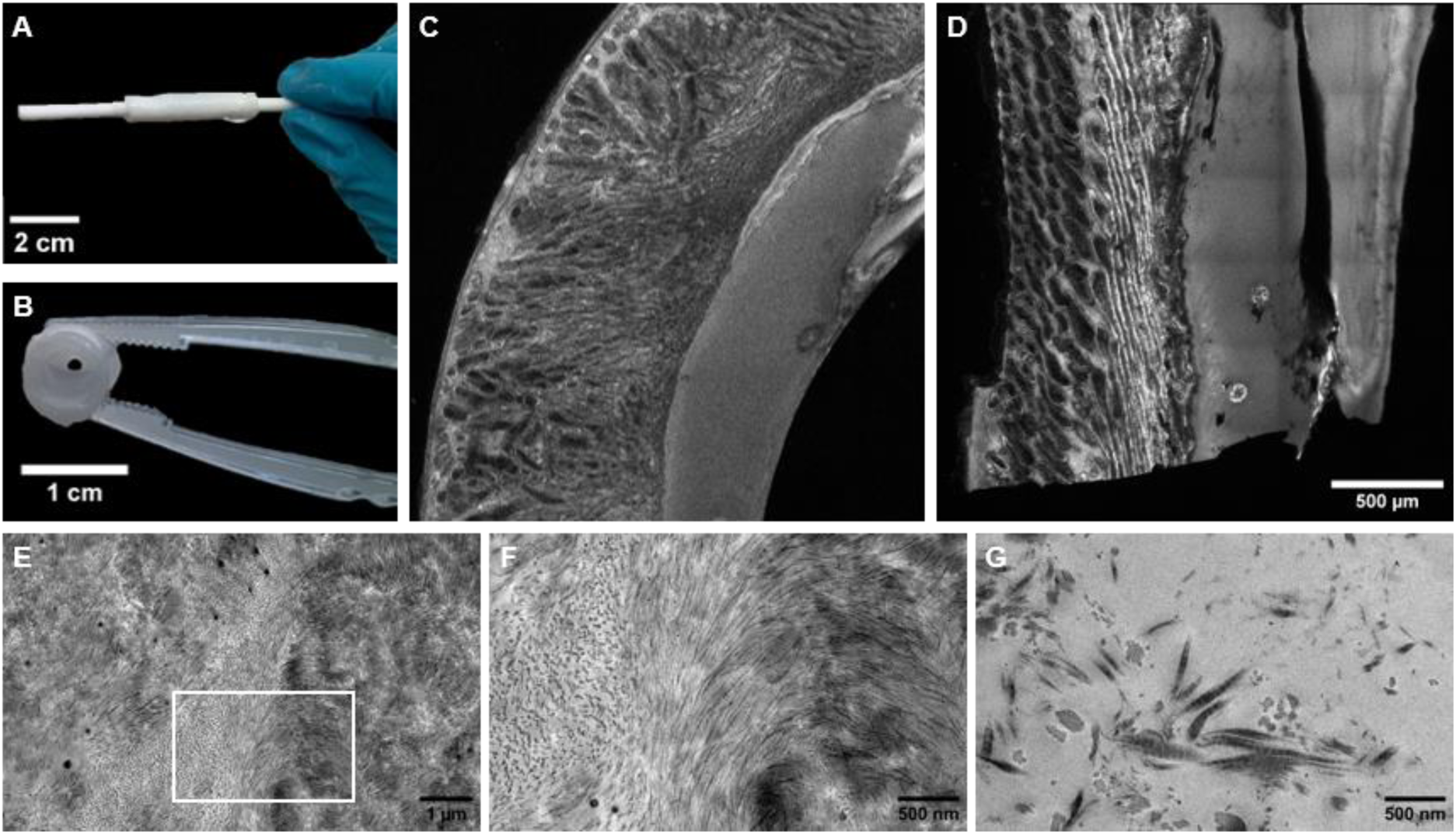
Bilayered collagen-based materials. A-B) Pictures of the obtained materials as observed along the long axis (A) or transversally (B). C-D) Confocal microscopy images of the transversal (C) and longitudinal (C) sections. E-G) TEM images on transversal sections of the scaffold displaying different sizes and densities of fibers. F) Higher magnification image of a specific part of the area imaged in E).

### 2.2. Mechanical characterization of collagen materials in comparison to native arteries

We assessed the mechanical behavior of the collagen tubes under loading conditions relevant to vascular tissues. *In vivo,* arteries experience two main types of deformation: radial expansion driven by pulsatile blood pressure and longitudinal stretching imposed by surrounding tissues and typically ranging from 15% to 20% in humans.^[39]^ To capture these two modes of deformation, mechanical characterization focused on arterial compliance, which quantifies radial deformation in response to physiological pressure variations between diastole and systole,^[40]^ and on the longitudinal Young’s modulus measured under uniaxial tensile loading, reflecting resistance to axial stretching.

A custom experimental setup was designed to independently probe radial and longitudinal mechanical responses (Figure 3A). Radial deformation was controlled by regulating the internal pressure using a syringe pump coupled to a pressure sensor, enabling precise compliance measurements. Longitudinal mechanical properties were assessed by applying uniaxial tensile loading along the tube axis using a motorized translation stage coupled to a force sensor. Measurements were performed on bilayered collagen tubes and native piglet carotid arteries, as well as on monolayer non-porous collagen tubes and monolayer porous collagen tubes.^[36]^ Bilayered materials appeared to be less brittle and more flexible than their non-porous collagen counterparts and could be clamped and maintained sealed at each extremity under normo- and hypertensive conditions. Monolayer smooth and porous collagen tubes could not be reliably evaluated under these conditions due to distinct failure mechanisms. Non-porous tubes tore during clamping, while porous tubes exhibited pressure-induced leakage caused by enlargement of the pore network, preventing stable pressurization beyond the hypotensive range. The watertightness of the scaffold was further confirmed by conducting tests using food dye injected into the lumen. Bilayered collagen tubes remained sealed (watertight) at internal pressures exceeding 200 mmHg, whereas leakage through monolayer porous tubes was observed below 100 mmHg (Supplementary Movies S1–S2).

**Figure 3.**
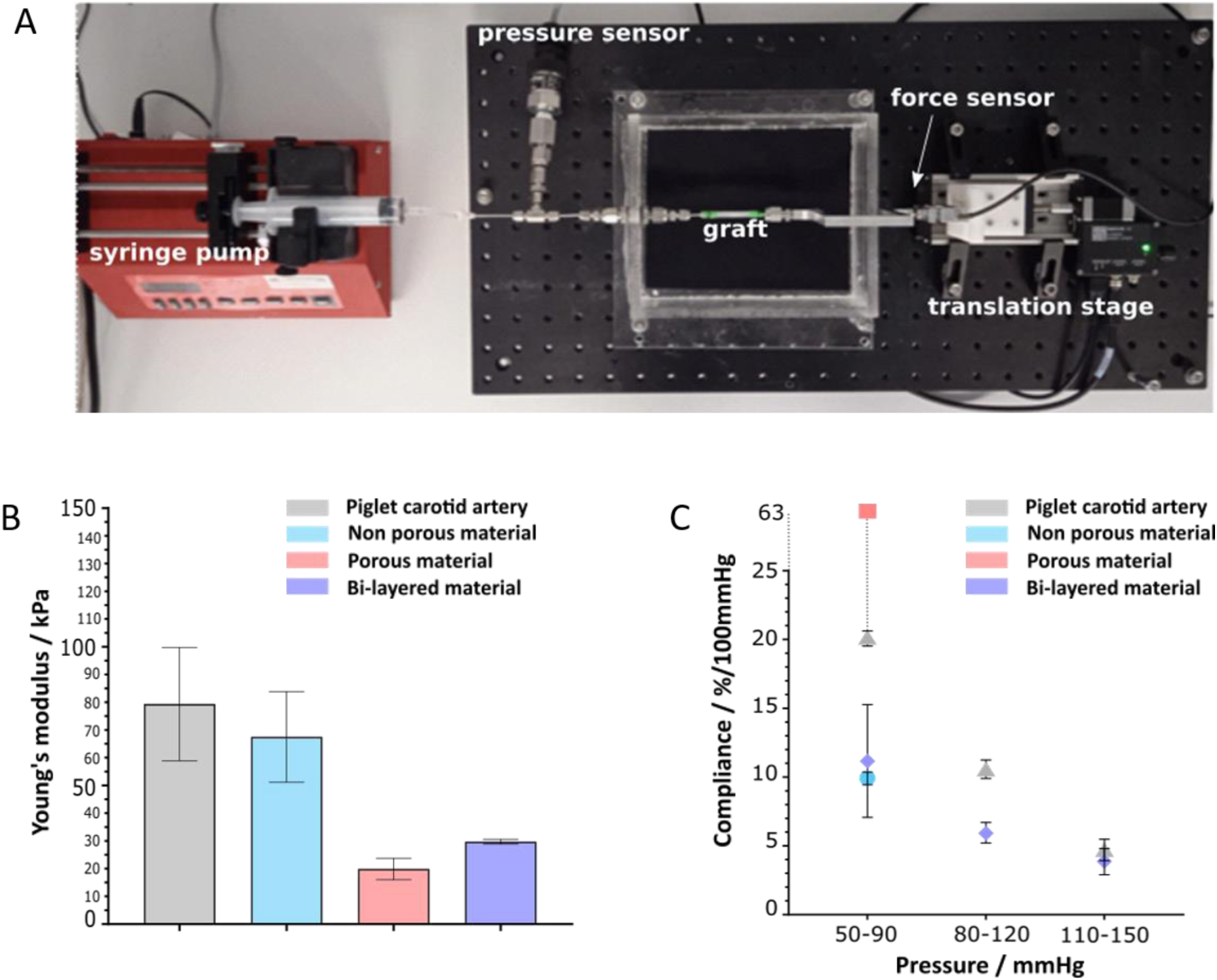
Mechanical evaluation of native piglet carotid artery and collagen-based materials. A) Home-made platform designed to enable control of applied hydrostatic pressure, of longitudinal elongation in a tubular configuration, as well as monitoring of radial deformation and applied force. B) Longitudinal tubular Young’s moduli values for each type of sample. N≥3 except for the porous materials, n=2. Error bars represent standard deviations. C) Compliance values for each type of sample, measured by analyzing radial deformation between two indicated values of pressures in the hypotensive, normotensive and hypertensive regimes. Only piglet carotid arteries and bilayered materials are shown at the 2 higher pressure ranges because of the failure of the porous and non porous collagen materials. N=3 under each condition except for the porous materials (n=1 in the hypotensive regime only) and the bilayered materials in the normo and hypertensive regimes (n=2). Error bars represent standard deviations.

Longitudinal tensile testing (Supplementary Movies S3–S6) showed that native piglet carotid arteries exhibited the highest Young’s modulus (79 ± 20 kPa, Figure 3B), followed by monolayer smooth collagen tubes (67 ± 16 kPa). Monolayer porous collagen tubes displayed substantially lower axial stiffness (20 ± 4 kPa), consistent with their highly porous architecture. Bilayered collagen tubes exhibited an intermediate Young’s modulus of 30 ± 1 kPa. This intermediate stiffness reflects the combined mechanical contributions of the dense luminal layer and the porous outer layer, yielding axial properties closer to those of native arteries than those of the porous configuration alone. In porous collagen materials, axial deformation is initially dominated by geometric rearrangement of the pore network rather than by direct stretching of collagen fibrils. The introduction of a dense, non-porous collagen layer promotes earlier load bearing by the fibrillar network, thereby shifting the mechanical response toward that of native arterial tissue. The axial stiffness values obtained for native arteries and monolayer smooth collagen tubes are consistent with previously reported measurements for uncrosslinked densified collagen tubular grafts and compacted tubular collagen gel constructs embedded with rat aortic smooth muscle cells.^[12,20]^ Moreover, longitudinal uniaxial testing of native human and porcine small-caliber arteries *ex vivo* has reported elastic moduli ranging from several tens to approximately 100 kPa.^[41,42]^ The axial stiffness measured for the bilayered constructs falls within the lower physiological regime of native artery mechanics, supporting their relevance as mechanically compatible vascular grafts.

Radial mechanical behavior was assessed through compliance measurements under pulsatile pressurization in the hypotensive (50–90 mmHg), normotensive (80–120 mmHg), and hypertensive (110–150 mmHg) regimes (Supplementary Movies S7–S10, Figure 3C), with all samples subjected to a physiological longitudinal pre-stretch of 20%. Porous and non-porous collagen materials could only be tested in the hypotensive regime due to mechanical failure at higher pressure. Monolayer smooth collagen tubes exhibited a compliance of 10 ± 0.5 %/100 mmHg, whereas monolayer porous tubes displayed markedly higher compliance values (63 /100 mmHg) (Figure 3-C). Bilayered collagen tubes and native piglet carotid arteries exhibited compliance values of 11 ± 4 %/100 mmHg and 20 ± 0.5 %/100 mmHg, respectively. With increasing pressure, both bilayered constructs and native arteries showed a pronounced reduction in compliance, reaching 6 ± 0.7 % and 10 ± 1.2 % under normotensive conditions, and 3.8 ± 1 % and 4.4 ± 1 % under hypertensive conditions, respectively. These values are consistent with those reported for native mammalian arteries and tissue-engineered blood vessels. In particular, König *et al.* reported comparable compliance values for native internal mammary arteries^[43]^ and showed that tissue-engineered blood vessels derived from adult human fibroblasts exhibited increasing compliance over time, reaching approximately 3.4 and 8.8 %/100 mmHg six months post-implantation.^[43,44]^ Dried collagen gels combined with elastin presented compliance values around 2.8%/100 mmHg, below those obtained here.^[17]^ Importantly, both the bilayered constructs and the native arteries exhibited a progressive decrease in compliance with increasing pressure regimes, reflecting the nonlinear elastic behavior characteristic of native blood vessels. This non-linear, strain-stiffening mechanical response observed for native arteries and some of the collagen materials arise from the intrinsic behavior of fibrillar collagen networks^[45,46]^ and are characteristic of native arterial tissues, whose mechanical response is largely governed by collagen organization and load-dependent fiber engagement.^[47,48]^ In arteries, strain stiffening is essential to accommodate large physiological variations in blood pressure and flow while preventing excessive wall deformation at high strain. These considerations are of particular interest for fully collagenous grafts capable of remodeling, as in the present study, and whose mechanical properties are expected to evolve after implantation through cellular colonization and ECM remodeling.^[44,49]^

### 2.3. In vitro cellularization of biomimetic tubular materials

#### HUVEC colonization on the luminal side of flat scaffolds in static conditions

As depicted in Supplementary Figure S3, HUVECs seeded on flat sections of the collagen scaffolds exhibited good spreading by day 5, and cell coverage was largely stable until day 14. VE-cadherin staining suggested the formation of a joint endothelial monolayer on the luminal surface.

#### HUVEC colonization in tubes under flow

HUVECs seeded on the luminal side of the tubular collagen scaffolds were subjected in a recirculating flow chamber to a steady shear stress of 1 dyn/cm² (0.1 Pa) for 3 or 5 days. Endothelialized matrix scaffolds maintained under static conditions were used as controls. At the end of the flow periods, samples were fixed and stained for actin (phalloidin) and nuclei (DAPI). Under static conditions, HUVECs formed a well-defined monolayer on the collagen substrate after 3 days of culture and exhibited sustained proliferation up to 5 days, resulting in a significant increase in cell density (p<0.05). In contrast, cells subjected to flow tended to show some albeit not statistically significant reduction in cell density, suggesting that flow may have led to limited impairment of cell proliferation and/or some cell detachment (Figure 4a). Analysis of actin staining revealed pronounced alignment in the direction of flow (Figure 4b). Flow also induced a certain level of nuclear elongation and prominent reorientation of the nuclear major axis in the direction of flow at the two time points studied (Figure 4c). These results demonstrate the ability to impose fluid dynamic shear stress in the tubular scaffolds and show that HUVECs cultured on the luminal side of these scaffolds respond to the applied shear stress.

**Figure 4.**
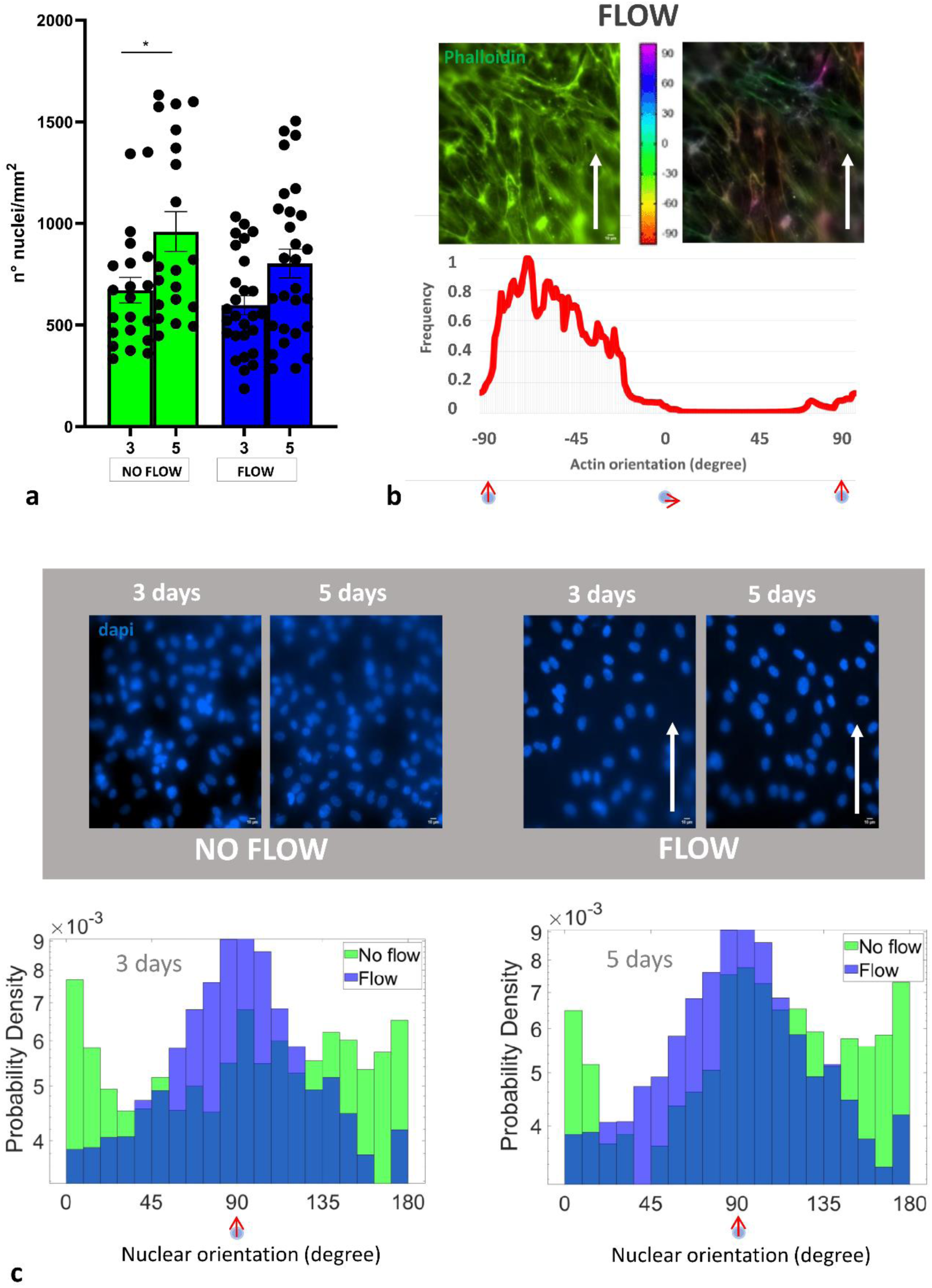
Endothelialization of the luminal side of the collagen matrix under static and flow (steady shear stress of 1 dyn/cm^2^) conditions for 3 and 5 days. (a) Quantification of cell proliferation based on the number of nuclei per unit area. (b) Distribution of actin orientation after 5 days of flow based on phalloidin staining. The distribution shows a clear peak near an angle of 0° (flow direction), indicating flow-induced actin alignment. (c) Histograms of nuclear orientation under static and flow conditions at the 3- and 5-day time points. Flow induces a shift in the distributions towards 0° (flow direction), indicating flow-induced nuclear alignment.

#### hMSC colonization on the adventitial (porous) side

Scaffolds were seeded on their adventitial (porous) side with 5x10^4^ or 10x10^4^ hMSCs and cultured up to 32 days. Samples were fixed at different time points, starting at 1 day, for histological analysis (Figure 5). hMSCs exhibited good adhesion to the scaffold surface at day 1 and initiated matrix colonization around day 5. hMSCs progressively penetrated the porous regions of the matrix, and colonization was obvious around day 10 to 12. This is supported by expression of metalloproteinase MMP9, suggesting that hMSCs remodel the scaffold while migrating (Figure 5).

**Figure 5.**
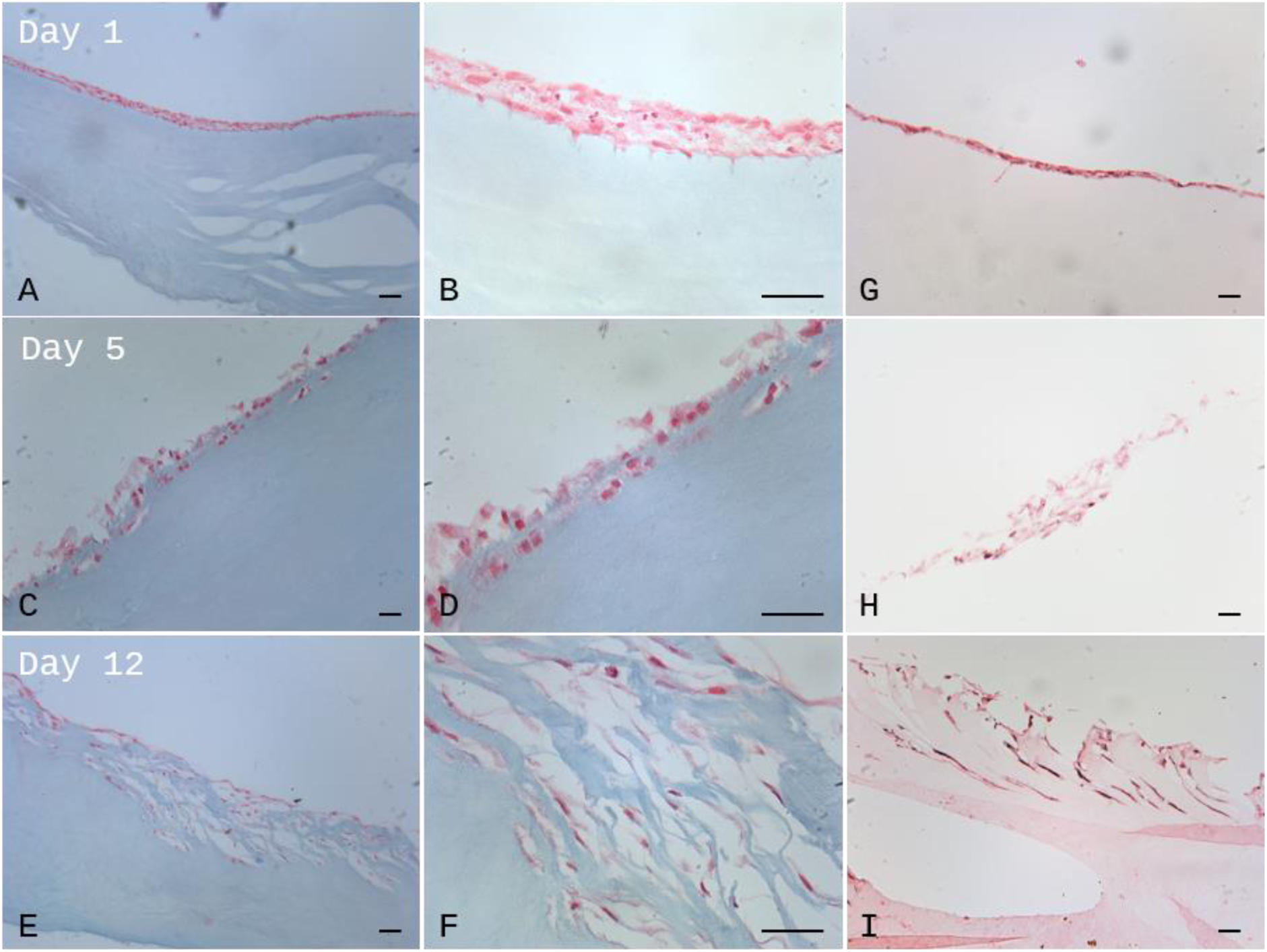
Colonization of the adventitial part by hMSCs. Samples were fixed at day 1 (A, B, G), day 5 (C, D, H), and day 12 (E, F, I) after seeding and stained with hematoxilin/eosin (A – F), or for MMP9 (G – I). Observation was conducted under 100X (A, C, E, G, H, I) and 400X magnification (B, D, F). Scale bars represent 50 µm.

After two weeks of culture, colonized scaffolds were exposed for two weeks either to hypoxia (3% O₂) or normoxia (21% O₂) in the presence or absence of TGF-β3, experimental conditions known to stimulate hMSC chondrogenic differentiation. No apoptotic cells were detected using TUNEL approach (data not shown). IHC staining for Sox9, a key chondrogenic transcription factor, showed that its expression was stimulated in response to hypoxia and to TGF-β3 (Figure 6 A-D). This suggests that hMSCs colonizing the scaffold remain able to respond to cytokines. In addition, areas colonized by hMSCs were positive for Alcian blue (Figure 6E-F), which allows the detection of extracellular matrix rich in glycosaminoglycans (GAGs). This supports the notion that hMSCs remodel the matrix and may secrete factors to form a pro-chondrogenic microenvironment.

**Figure 6.**
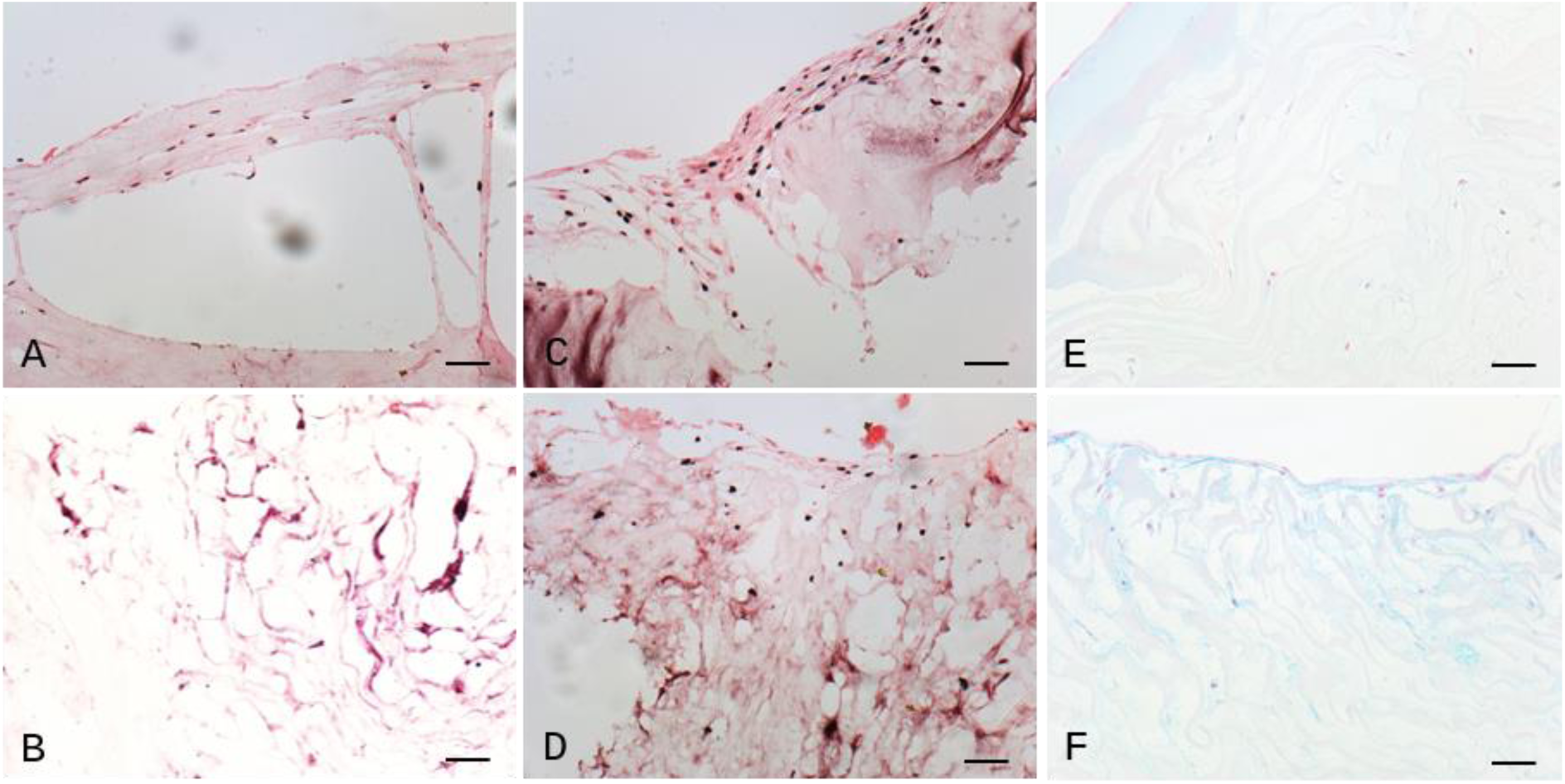
hMSC colonization after 4 weeks in culture, with 2 first weeks under normoxia conditions. Samples were fixed after the 2 last weeks of culture under normoxia (A, C, E), hypoxia (B, D, F), in the absence (A, B) or the presence of TGF-β3 (C – F), and stained for Sox9 (A – D) or colored with Alcian blue (E & F). Observation was conducted under 200X magnification. Scale bars represent 50 µm.

### 2.4 Surgical feasibility study

Surgical feasibility was investigated based on manipulation of the materials by surgeons, including *ex vivo* anastomosis with calf bronchus segment, along with quantitative assessment of suture retention strength (SRS) in comparison to that in native arteries. Before suturing, the scaffold qualitatively displayed adequate elasticity and cohesive handling properties.

To evaluate the suitability of the collagen tubular scaffold for tracheal substitution, two complementary suturing tests were performed reproducing key steps of airway reconstruction, following previously reported procedures.^[6–8]^. Both end-to-end anastomosis and longitudinal wall suturing were assessed, with silicone stent insertion to mimic the temporary stenting phase required in clinical airway bioengineering. The stent serves to prevent early collapse of the graft, as the scaffold does not initially possess the radial force of native airways.

Suture retention strength is defined as the force necessary to pull a suture from a material, here corresponding to the maximum traction force applied by means of the custom platform on a thread before tearing the material (Supplementary Figure S4). Measurements performed on piglet carotid arteries and bilayered collagen materials revealed that SRS in the scaffolds is ten times lower than that in native tissues, with values of 0.15 ± 0.01 (n=4) and 2 ± 0.15 N (n=2), respectively (Supplementary Figure S4). Suture retention strength of tissue-engineered blood vessels derived from adult human fibroblasts was reported to be not significantly different from the value measured on internal mammalian artery, around 1.4 N (152 ± 50 versus 138 ± 50 gF, respectively).^[43]^ Still, tubular conduits obtained from collagen gels and elastin-like protein polymers that could further be implanted in rats displayed values of SRS around 300 mN (30.5 ± 4.2 and 37.0 ± 5.1 gF depending on diameter).^[17]^ Furthermore, dehydrated and crosslinked 0.5 mm-diameter dense collagen tubes having SRS values of 7.9 ± 2.2 gF were implanted as interpositional grafts in the rat femoral circulation with stable anastomosis for 20 min.^[15]^

As previously mentioned, before suturing the scaffold exhibited adequate elasticity and cohesive handling properties, but during a first suturing test a degree of fragility became evident once the needle and Prolene 6/0 thread were passed through the wall, with a tendency towards local tearing at the puncture sites during suture tightening. Despite this limitation, it was possible to exert tension on thread sutured at the extremity of the material (Movie S11), and both end-to-end and longitudinal anastomoses could be completed when performed with careful tension control (Figure 7A).

**Figure 7.**
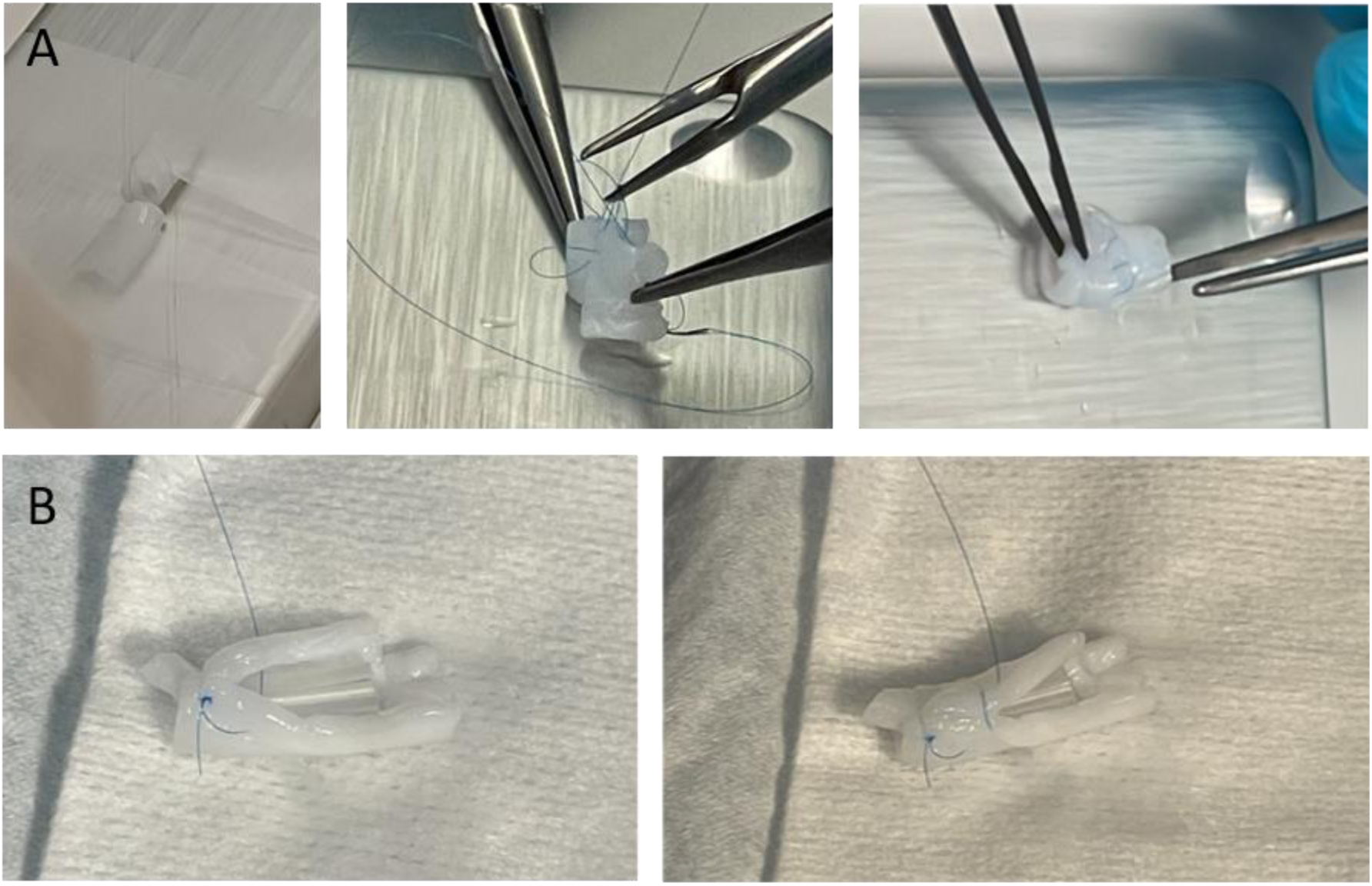
A) End-to-end Prolene suture of the scaffold. B) Longitudinal suture with internal silicone stent.

A second suturing test using Monocryl 6/0 thread to perform scaffold–bronchus anastomosis confirmed the overall mechanical response of the scaffold during manipulation: good flexibility before suturing but noticeable fragility during needle passage and traction. Nevertheless, the anastomosis could be completed without disruption of the material, yielding a technically satisfactory and promising result (Supplementary Movie S12 and Figure 8). The anastomosis was reinforced with TachoSil**®**, a fibrin-coated collagen patch widely used in thoracic and abdominal surgery to enhance sealing reinforcement, which contributed to improved stability of the suture line.

**Figure 8:**
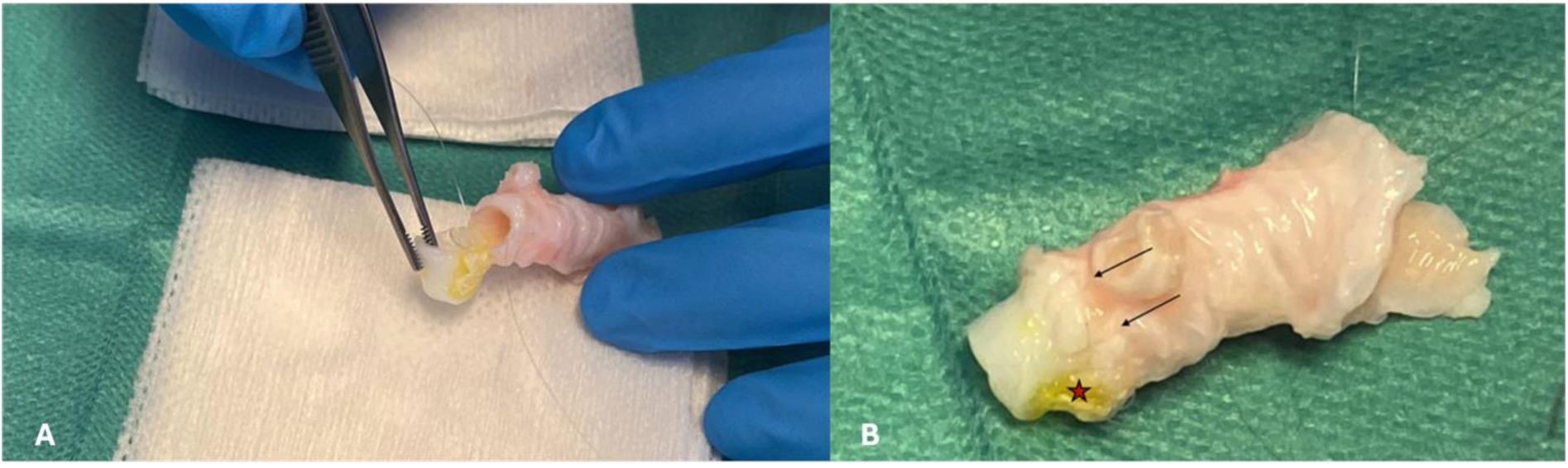
A) Scaffold–bronchus anastomosis reinforced with TachoSil - B) Completed anastomosis (arrows: suture line; star: TachoSil).

## 3. Conclusion

The tubular materials (4 and 8 mm internal and external diameters) presented here consist purely of type I collagen and the process of fabrication enables topological control over the porosity, with successful fabrication of a tunable bilayered collagen tube combining a dense, smooth luminal surface with a radially tunneled porous outer layer, seamlessly integrated within a single continuous material. Hybrid functional properties are achieved, with radially oriented tunnels of the outer layer providing an architecture favorable to cell colonization from adjacent tissues and nutrient transport, while the dense and smooth luminal surface promotes endothelialization and tube tightness to withstand physiological and supraphysiological pressurization, and ensuring lumen sealing. This combination therefore provides mechanically and functionally relevant collagen-based scaffolds for vascular and airway applications. In particular, the obtained tubes promoted formation of a joint endothelium with recapitulation of physiological endothelial cell orientation under flow. Colonizing human mesenchymal stem cells remodeled the matrix and under appropriate conditions may secrete factors to form a pro-chondrogenic microenvironment. Suturing tests also demonstrated that the scaffolds can be handled, incised, sutured and stented similarly to clinical airway substitutes, that end-to-end and longitudinal sutures are feasible despite localized fragility, and that the material presents promising surgical manipulability. The sequential fabrication steps enable precise and reproducible control over the final tube dimensions, including outer diameter, inner diameter, and the relative thicknesses of the porous and dense layers. As a result, the fabrication strategy is readily adaptable to the geometrical requirements of different tubular biological tissues. By providing on-demand tissues with precisely modulated properties, the development of such tubular biomaterials represents a promising alternative for the treatment of multiple diseases.

## 4. Experimental Section/Methods

### Fabrication of bilayered collagen scaffolds

Collagen I was extracted from young rat tail tendons. After thorough cleaning of tendons with phosphate buffered saline (PBS; 1X) and 4 M NaCl, tendons were dissolved in 3 mM HCl. Differential precipitation with 300 mM NaCl and 600 mM NaCl, followed by redissolution and dialysis in 3 mM HCl, provided collagen of high purity. The final collagen concentration was determined using hydroxyproline titration with a spectrophotometer. To attain a high fibrillar collagen density in the model, the remaining collagen solutions were concentrated at 40 mg.mL^-1^. The solutions were then transferred into Vivaspin tubes with 300 kDa filter and centrifuged at 3000 g at 10°C, to reach this final concentration.

Concentrated collagen at 40 mg.mL^-1^ was introduced in between two cylindrical molds (inner aluminum mold: 5 mm; outer PTFE mold : 8 mm), with the top end hermetically sealed by a cap and a remaining hole at the bottom to let the aluminum mold outside. For removal of air bubbles, the sample was centrifuged at 1g at 10°C for 10 minutes. The sample was further placed onto a home-made set-up that allowed continuous dipping of the tip of the aluminum mold in liquid nitrogen for 10 min.^[36]^ After this ice templating step, the metallic mold and the caps were removed from the frozen sample, which was further exposed to ammonia vapors at 0°C for 48h to induce the topotactic fibrillogenesis. The sample was then placed for 24 h in water vapor in a heat room at 37°C for removal of ammonia vapors. The first porous layer of fibrillated collagen was washed 3 times in PBS (1X) for 15 min. Acidic collagen solution at 26 mg.mL^-1^ was added one-third at a time inside the tubular scaffold (4 mm) and centrifuged at 1g at 10°C for 5 min between each each collagen addition step. A PTFE rod of 4 mm was introduced inside the tube and sealed on one side by a regular cap and the other side by a cap with holes to allow buffer diffusion. The whole sample was recentrifuged at 1g at 10°C for 5 min and left at room temperature for 45 min to allow dissolution of fibrillated collagen to mix with newly non fibrillated collagen. It was then immersed in PBS (5X) for 1 h. The caps were removed and the sample immersed again in PBS (5X), and the fibrillogenesis was completed by keeping the sample in PBS (5X: 685 mM NaCl, 13.4 mM KCl, 40.35 mM Na_2_HPO_4_, and 7.35 mM NaH_2_PO_4_) for 14 days at room temperature.

### Confocal microscopy

After fibrillogenesis, 200 µm-thick sections were cut. Collagen was labeled with fluorescein isothiocyanate (FITC, 0.05 mg/mg of collagen) at 4°C overnight. Observations were conducted using a Zeiss LSM 980 upright confocal microscope.

### Transmission electron microscopy

Hydrated samples were cross-linked with 2.5% paraformaldehyde (PFA), 2% glutaraldehyde, 0.18M sucrose and 0.1% picric acid for 12 h. The samples were subsequently post-fixed with uranyl acetate in ethanol for 12 h and dehydrated using baths with increasing concentrations of ethanol. Samples were incorporated in SPURR-S resins prior to sectioning. 70 nm ultrathin sections (Leica microtome) were contrasted with uranyl acetate and observed on a transmission electron microscope (FEI Tecnai Spirit G2) operating at 120 kV to observe the ultrastructural collagen features. Images were recorded on a Gatan CCD camera.

### Porosity quantification

Porosity on transversal section of fibrillated materials was quantified using ImageJ Fiji for analyzing confocal microscopy images. Images were first pseudo-binarized using the LabKit segmentation plugin and porosity was quantified using the RidgeDetection plugin. This latter detects ridges or linear structures in images by analyzing intensity variations and identifying pixels with high gradient values along a specific direction. In the end, the mean width of the pores and pore walls was retrieved. Pore surfacic porous fraction was measured by dividing the area of pores by the total area.

### Harvesting of native artery and bronchus specimens

Calf bronchi and piglet carotid arteries were recovered immediately after animal’s sacrifice from waste produced during educational sessions at the hospital.

### Fabrication of porous and non-porous collagen materials

Concentrated collagen is introduced in between two cylindrical molds (ABS insulating inner mold: 3 × 5 mm; aluminum conductive outer mold: 8 × 10 mm: length: 50 mm) following the previously published protocol.^[36]^ The sample is placed onto the home-made set-up that allows a continuous and regulated dipping of the sample in liquid nitrogen. Once frozen, the outer mold is quickly removed and two fibrillogenesis conditions are applied. Porous materials result from exposure to ammonia vapors, with subsequent distilled water vapors exposure, as previously described, while non porous materials result from exposure to PBS bath (10×: 1.37 M, KCl 26.8 mM, Na2HPO4 80.7 mM, and NaH2PO4 14.7 mM) at −3 °C in a thermostatic bath for 72 h. For both types of materials, fibrillogenesis is completed by keeping the sample in PBS (5X) for 14 days at room temperature, as previously described.

### Mechanical measurements

A custom-made setup was used, which can be divided into two main components (Figure 3-A). First, a pressure sensor (Trafag 8287) was connected to a syringe pump (KF Technology) to provide feedback control, enabling precise regulation of the pressure applied to the tubular sample, thereby enabling compliance measurements. Second, a translation stage (Zaber, X-LHM050A) linked to a force sensor (ME-Meßsysteme GmbH, Germany) was employed to measure the force required to deform the sample. Each end of the tubular materials was sealed, either with a solid cap at the distal end or with a perforated cap at the proximal end (i.e., the part of the tube connected to the millifluidic device). All measurements were conducted under immersion in 1X PBS at 37°C.

#### Tubular traction force and Suture Retention Strength measurement

Uniaxial tensile testing of tubular samples was conducted using a custom-made stage equipped with a 10 N load cell (ME-Meßsysteme GmbH, Germany). The initial distance between the caps was adjusted until each tube was extended with a measurable non-zero force, defining this distance as the sample’s unloaded length. To ensure consistency and accuracy, each tube underwent two pre-conditioning cycles to produce a stable force-elongation profile. Cyclic testing was then carried out on each tube, with data from the third cycle’s loading phase analyzed. The deformation rate, defined as 𝑘 = 𝑣 / 𝐿, where 𝑣 is the deformation speed and 𝐿 is the unloaded sample length, was maintained at a constant value of 0.8% for all samples. This protocol was uniformly applied across all tests. The resulting stress-strain curves were used to calculate the Young’s modulus, focusing exclusively on the linear portion of these curves, corresponding to 10-20% strain. At least three tubular samples were tested under each condition except for the porous materials (2 samples).

For suture retention strength measurements, the same uniaxial tensile testing setup described above was used. Native arteries or bilayered collagen constructs were longitudinally opened and cut into rectangular flat specimens (30 mm × 10 mm). A Prolene 5/0 suture was placed at the opposite end, 3 mm from the free edge and centered across the width, to prevent premature edge failure and ensure uniform stress distribution. Tensile loading was applied along the longitudinal axis of the specimen at 0.1 mm·s⁻¹ until failure. Force was recorded over time, and the maximum force prior to failure was defined as the suture retention strength. Representative force–time curves are provided in S4.

#### Compliance measurement

Compliance was calculated using the formula:

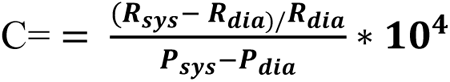

where 𝑹_𝒔𝒚𝒔_ and 𝑹_𝒅𝒊𝒂_ are the systolic and diastolic internal radii, respectively, and 𝑷_𝒔𝒚𝒔_ and 𝑷_𝒅𝒊𝒂_ are the systolic and diastolic pressures, respectively.^[40]^ Compliance values were measured under different pulsatile regimes: hypotensive (50-90 mmHg), normotensive (80-120 mmHg), and hypertensive (110-150 mmHg). The measurements were conducted on samples pre-stretched to 20%, reflecting the average physiological pre-stretch values of arteries reported in the literature.^[39]^ Additionally, we measured the pre-stretch values of native arteries under free inflation to 100 mmHg and found similar pre-stretch values (data not shown). The pressure oscillation between systolic and diastolic values was maintained at approximately 1 Hz. Each compliance value for each sample was measured three times for each pulsatile regime. The tube diameter at the center of the region of interest was measured as a function of the applied systolic and diastolic pressures using a CCD camera. For each pressure condition, the diameter was determined by manual image analysis using ImageJ. At least three tubular samples were tested under each condition except for the porous materials (1 sample) in the hypotensive regime and the bilayered materials in the normo and hypertensive regimes (2 samples).

### Cellular colonization

#### HUVECs under static conditions

Human umbilical vein endothelial cells (HUVECs, Lonza) in passages 5-8 were cultured at 37°C and 5% CO_2_ in EGM2 cell medium (Lonza). HUVECs were seeded on the luminal surface of 6 mm-diameter discs of the collagen scaffolds at a density of 5,000 cells/mm^2^ and cultured for 14 days. At the end of the culture period, samples were fixed with 4% paraformaldehyde and permeabilized with 0.1% Triton X-100. Samples were incubated with primary VE-cadherin antibody (Anti-VE-Cadherin Antibody, clone Vli37, Merck) for 1 hour at room temperature. A mixture of Alexa-fluor phalloidin 488 (Invitrogen™, Thermo Fisher Scientific) at 1/2000 and secondary antibody (Donkey Anti-Rabbit IgG H&L (Alexa Fluor® 647, Abcam) diluted at 1/400 in PBS was further added for 1 hour at room temperature. A series of z-stack images were acquired using a Zeiss LSM 980 upright confocal microscope.

#### HUVECs under flow conditions

HUVECs were seeded on bilayered collagen tubes at a density of 1,000 cells/mm^2^. After cell adhesion, the tubes were inserted in a recirculating flow loop and subjected to a steady shear stress of 1 dyn/cm^2^ (0.1 Pa) for a period of 3 or 5 days. Cells seeded on similar collagen tubes and not subjected to flow were used as controls.

##### Immunofluorescence staining and microscopy

Samples were fixed with 4% paraformaldehyde, permeabilized with 0.1% Triton x-100, and then incubated with DAPI (Sigma Aldrich, ref. D1542) and phalloidin 495 (LifeTechnologies, ref. A12381) to visualize the nuclei and the F-actin cytoskeleton, respectively. A series of images in the z-direction (z-stacks) were acquired at 1 µm intervals. Cell images were acquired using an inverted microscope (Nikon Eclipse Ti-U, Japan) equipped with a CCD camera (Orca AG; Hamamatsu, Japan) and a motorized x-y stage equipped with a 20X objective.

##### Analysis of immunofluorescence microscopy images

Cell proliferation was quantified by counting the number of nuclei based on the DAPI signal using the Cell Counter plugin in FIJI. Nuclear shape was analyzed based on circularity values obtained in FIJI from the DAPI images. Nuclear orientation was defined as the angle between the major cell axis and the flow direction. Actin orientation was determined using the Orientation J plugin in FIJI.

##### Statistical analysis

Data were expressed as mean ± SE (standard error). Statistical analyses were performed in GraphPad Prism (GraphPad Software) using the Mann-Whitney *t* test with p < 0.05 considered statistically significant.

### External surface seeding with human mesenchymal stem cells (hMSC)

Six mm-diameter discs were seeded on their external surface with 50,000 or 100,000 hMSCs and cultured up to 4 weeks in DMEM containing 25 mM D-glucose, 10 mM HEPES, 23.8 mM NaHCO3, 1 mM sodium pyruvate, 2 mM L-glutamine, 100 U/ml penicillin, 100 µg/ml streptomycin, 10 µg/ml gentamycin, and supplemented with 10% FCS, in a 5% CO_2_–95% air atmosphere. After two weeks of culture, the medium was supplemented with 1% dexamethasone, 0.1% ascorbic acid, 0.1% L-proline, 1% Insulin-Transferrin-Selenium mix, and samples were exposed either to hypoxia (3% O₂) in a hypoxia workstation (InVivo2 300, Baker Ruskinn) or to normoxia (21% O₂), in the presence or absence of 10 ng/mL TGF-β3 (R&D systems Biotechne) for two more weeks. The medium was changed every 48 h.

#### Histological treatment and immuno histochemical staining

At different time points, samples were fixed with 4% paraformaldehyde in PBS (1×) for 16 h at 4 °C, then rinsed in PBS. Samples were successively transferred in 30%, 50% and 70% ethanol bath for dehydration and finally embedded in paraffin in order to prepare 5 µm-thick sections (TisCell13 histology facility, UFR SMBH). Sections were stained with haematoxylin and eosin dyes or with alcian blue pH 2.5 dye, or stained for MMP9 (Abcam#137867) or Sox9 (Abcam#185230). Antigen retrieval was performed in a boiling Tris-EDTA buffer (10 mM Tris, 1.2 mM EDTA, pH 9.0). Endogenous peroxidases were quenched with 0.5% hydrogen peroxide in methanol for 15 min, and sections were incubated with 4% normal horse serum for 1 h to block nonspecific antibody binding sites. Sections were incubated with the different primary antibodies overnight at 4^◦^C. The next day, slides were rinsed with PBS/0.1% TritonX100 and incubated with biotinylated anti-rabbit secondary antibodies (Vector Laboratories) for 30 min. After washes, samples were incubated with peroxidase (Vectastain Elite ABC kit, Vector Laboratories) for 30 min. DAB (3,3’-diaminobenzidine) solution was used to visualize the positive reactions, and slides were counterstained with nuclear fast red (Sigma Aldrich).

### Suturing tests

End-to-end anastomoses were first created at both tubular extremities using continuous 6/0 Prolene sutures. A longitudinal incision was then performed, and a second continuous suture was completed after insertion of a silicone stent. To reproduce a more clinically realistic configuration, a continuous 6/0 Monocryl anastomosis was further performed between the scaffold and a calf bronchus segment. The suture line was reinforced using TachoSil®, a fibrin-coated collagen patch routinely applied to improve sealing and mechanical support in thoracic surgery.

## References

[1] I. Martinier, L. Trichet, F. M. Fernandes, Chem. Soc. Rev. 2025, 54, 790.

[2] M. Carrabba, P. Madeddu, Front. Bioeng. Biotechnol. 2018, 6, 41.

[3] E. S. Clark, C. Best, E. Onwuka, T. Sugiura, N. Mahler, B. Bolon, A. Niehaus, I. James, N. Hibino, T. Shinoka, J. Johnson, C. K. Breuer, Journal of Pediatric Surgery2016, 51, 49.

[4] H. Etienne, D. Fabre, A. Gomez Caro, F. Kolb, S. Mussot, O. Mercier, D. Mitilian, F. Stephan, E. Fadel, P. Dartevelle, EurRespir J2018, 51, 1702211.

[5] Y. Liang, S. Wei, A. Zhang, Regenerative Therapy 2025, 29, 364.

[6] E. Martinod, K. Chouahnia, D. M. Radu, P. Joudiou, Y. Uzunhan, M. Bensidhoum, A. M. Santos Portela, P. Guiraudet, M. Peretti, M.-D. Destable, A. Solis, S. Benachi, A. Fialaire-Legendre, H. Rouard, T. Collon, J. Piquet, S. Leroy, N. Vénissac, J. Santini, C. Tresallet, H. Dutau, G. Sebbane, Y. Cohen, S. Beloucif, A. C. d’Audiffret, H. Petite, D. Valeyre, A. Carpentier, E. Vicaut, JAMA2018, 319, 2212.

[7] E. Martinod, D. M. Radu, I. Onorati, A. M. S. Portela, M. Peretti, P. Guiraudet, M.-D. Destable, Y. Uzunhan, O. Freynet, K. Chouahnia, B. Duchemann, J. Kabbani, C. Maurer, P.-Y. Brillet, L. Fath, E. Brenet, C. Debry, C. Buffet, L. Leenhardt, D. Clero, N. Julien, N. Vénissac, F. Tronc, H. Dutau, C.-H. Marquette, C. Juvin, G. Lebreton, Y. Cohen, E. Zogheib, et al., American Journal of Transplantation 2022, 22, 2961.

[8] E. Martinod, D. M. Radu, I. Onorati, X. Chapalain, A. M. Santos Portela, M. Peretti, O. Freynet, Y. Uzunhan, K. Chouahnia, B. Duchemann, C. Juvin, G. Lebreton, H. Rouard, G. Van Der Meersch, G. Galvaing, J.-B. Chadeyras, F. Tronc, P. Kuczma, C. Trésallet, N. Vénissac, S. Beloucif, O. Huet, E. Vicaut, JAMA Surg 2025, 160, 912.

[9] L. Cen, W. Liu, L. Cui, W. Zhang, Y. Cao, Pediatr Res 2008, 63, 492.

[10] C. B. Weinberg, E. Bell, 231.

[11] J. Hirai, T. Matsuda, Cell Transplant 1996, 5, 93.

[12] C. L. Cummings, D. Gawlitta, R. M. Nerem, J. P. Stegemann, Biomaterials2004, 25, 3699.

[13] S. Meghezi, D. G. Seifu, N. Bono, L. Unsworth, K. Mequanint, D. Mantovani, JoVE 2015, 52812.

[14] D. G. Seifu, S. Meghezi, L. Unsworth, K. Mequanint, D. Mantovani, Journal of the Mechanical Behavior of Biomedical Materials 2018, 80, 155.

[15] X. Li, J. Xu, C. T. Nicolescu, J. T. Marinelli, J. Tien, 2017, 23, 335.

[16] C. E. Ghezzi, B. Marelli, N. Muja, S. N. Nazhat, 2012, 8, 1813.

[17] V. A. Kumar, J. M. Caves, C. A. Haller, E. Dai, L. Liu, S. Grainger, E. L. Chaikof, 2013, 9, 8067.

[18] C. Loy, A. Lainé, D. Mantovani, 2016, 11, 1673.

[19] C. Loy, D. Pezzoli, G. Candiani, D. Mantovani, 2018, 13, 1700359.

[20] A. W. Justin, F. Cammarata, A. A. Guy, S. R. Estevez, S. Burgess, H. Davaapil, A. Stavropoulou-Tatla, J. Ong, A. G. Jacob, K. Saeb-Parsy, S. Sinha, A. E. Markaki, 2023, 145, 213245.

[21] M. Madaghiele, A. Sannino, I. V. Yannas, M. Spector, J Biomedical Materials Res 2008, 85A, 757.

[22] K. M. Pawelec, R. J. Wardale, S. M. Best, R. E. Cameron, 2015, 26, 13.

[23] M. J. W. Koens, K. A. Faraj, R. G. Wismans, J. A. Van Der Vliet, A. G. Krasznai, V. M. J. I. Cuijpers, J. A. Jansen, W. F. Daamen, T. H. Van Kuppevelt, 2010, 6, 4666.

[24] Y. Xu, J. Dai, X. Zhu, R. Cao, N. Song, M. Liu, X. Liu, J. Zhu, F. Pan, L. Qin, G. Jiang, H. Wang, Y. Yang, 2022, 34, 2106755.

[25] C. O’Leary, B. Cavanagh, R. E. Unger, C. J. Kirkpatrick, S. O’Dea, F. J. O’Brien, S.-A. Cryan, 2016, 85, 111.

[26] T. Khalid, L. Soriano, M. Lemoine, S.-A. Cryan, F. J. O’Brien, C. O’Leary, Front. Bioeng. Biotechnol. 2023, 11, 1187500.

[27] L. Soriano, M. Lemoine, B. Cavanagh, A. Johnston, T. Khalid, F. J. O’Brien, C. O’Leary, S.-A. Cryan, ACS Biomater. Sci. Eng. 2025, 11, 5293.

[28] J. F. Cavallaro, P. D. Kemp, K. H. Kraus, Biotech & Bioengineering 1994, 43, 781.

[29] F. Zhang, Y. Xie, H. Celik, O. Akkus, S. H. Bernacki, M. W. King, 2019, 11, 035020.

[30] L. Magnan, G. Labrunie, M. Fénelon, N. Dusserre, M.-P. Foulc, M. Lafourcade, I. Svahn, E. Gontier, J. H. Vélez V., T. N. McAllister, N. L’Heureux, 2020, 105, 111.

[31] L. J. Chamberlain, I. V. Yannas, A. Arrizabalaga, H.-P. Hsu, T. V. Norregaard, M. Spector, 1998, 19, 1393.

[32] B. W. Riblett, N. L. Francis, M. A. Wheatley, U. G. K. Wegst, 2012, 22, 4920.

[33] M. C. T. Asuncion, J. C.-H. Goh, S.-L. Toh, 2016, 67, 646.

[34] C. Rieu, C. Parisi, G. Mosser, B. Haye, T. Coradin, F. M. Fernandes, L. Trichet, ACS Appl. Mater. Interfaces2019, 11, 14672.

[35] C. Parisi, B. Thiébot, G. Mosser, L. Trichet, P. Manivet, F. M. Fernandes, Biomater. Sci. 2022, 10, 6939.

[36] I. Martinier, F. Fage, A. Kakar, A. Castagnino, E. Saindoy, J. Frederick, I. Onorati, V. Besnard, A. I. Barakat, N. Dard, E. Martinod, C. Planes, L. Trichet, F. M. Fernandes, Biomater. Sci. 2024, 12, 3124.

[37] K. Yin, P. Divakar, U. G. K. Wegst, 2019, 84, 231.

[38] L. Besseau, M.-M. Giraud-Guille, 1995, 251, 197.

[39] R. A. Macrae, K. Miller, B. J. Doyle, 2016, 52, 380.

[40] D. B. Camasão, D. Mantovani, 2021, 10, 100106.

[41] L. Horny, T. Adamek, E. Gultova, R. Zitny, J. Vesely, H. Chlup, S. Konvickova, 2011, 4, 2128.

[42] G. Sommer, P. Regitnig, L. Költringer, G. A. Holzapfel, 2010, 298, H898.

[43] G. Konig, T. N. McAllister, N. Dusserre, S. A. Garrido, C. Iyican, A. Marini, A. Fiorillo, H. Avila, W. Wystrychowski, K. Zagalski, M. Maruszewski, A. L. Jones, L. Cierpka, L. M. De La Fuente, N. L’Heureux, 2009, 30, 1542.

[44] N. L’Heureux, N. Dusserre, G. Konig, B. Victor, P. Keire, T. N. Wight, N. A. F. Chronos, A. E. Kyles, C. R. Gregory, G. Hoyt, R. C. Robbins, T. N. McAllister, 2006, 12, 361.

[45] A. J. Licup, S. Münster, A. Sharma, M. Sheinman, L. M. Jawerth, B. Fabry, D. A. Weitz, F. C. MacKintosh, Proc. Natl. Acad. Sci. U.S.*A.* 2015, 112, 9573.

[46] D. Vader, A. Kabla, D. Weitz, L. Mahadevan, PLoS ONE 2009, ee5902

[47] J. E. Wagenseil, R. P. Mecham2009, 89, 957.

[48] G. Martufi, T. C. Gasser, 2011, 44, 2544.

[49] S. Pashneh-Tala, S. MacNeil, F. Claeyssens, 2016, 22, 68.

