## Supplemental information for "Collagen-based bilayered biomimetic tubular materials for vascular and airway applications"

### Collagen-based double layered biomimetic tubular materials for vascular and tracheal applications

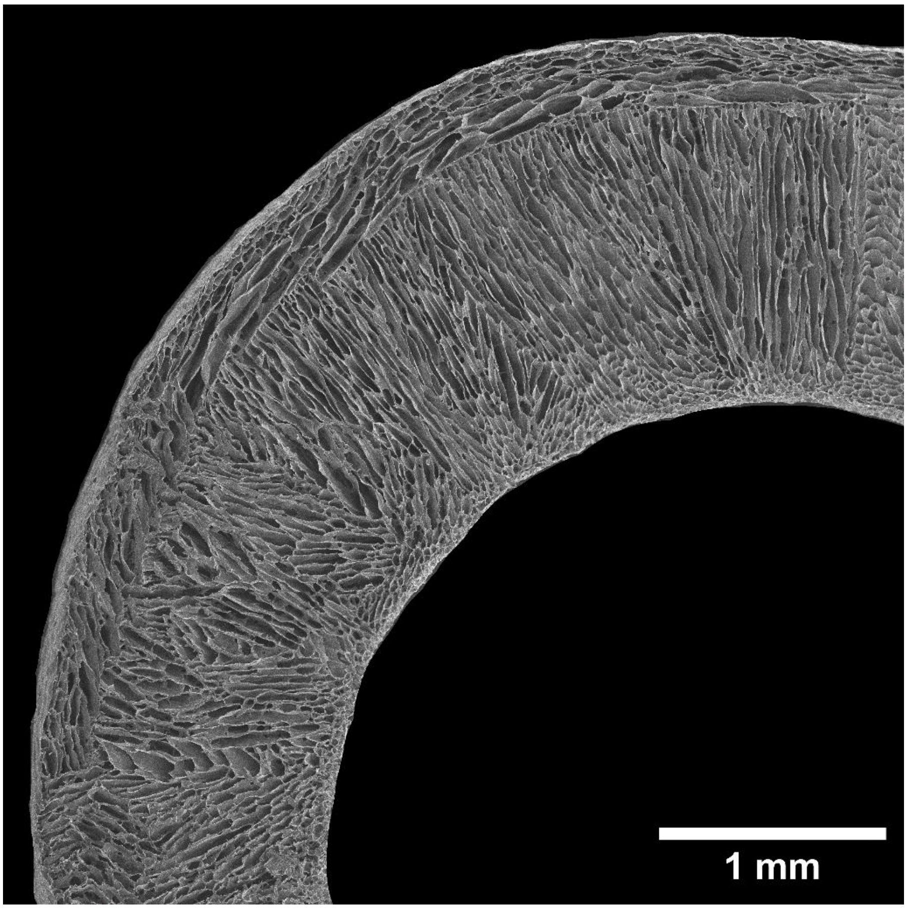


**Figure S1.** Reconstitution of the transversal section of the first layer of type I collagen material as obtained after the ice templating step and imaged by scanning electron microscopy after lyophilization.

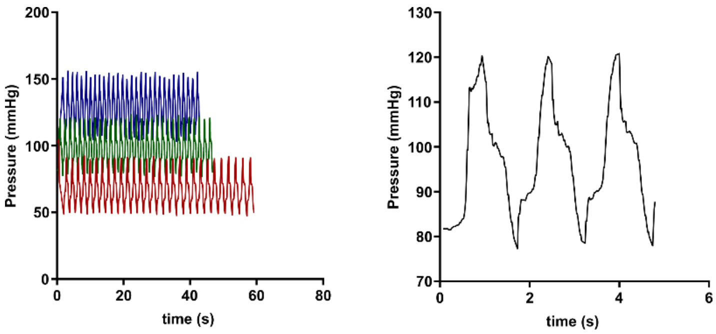


**Figure S2.** Inflation analysis of native piglet carotid artery in normo-, hypo- and hypertensive regimes, with magnification of the signal in the normotensive regime (right). The pressure oscillation frequency between the systolic and diastolic phases was maintained at approximately 1 Hz.

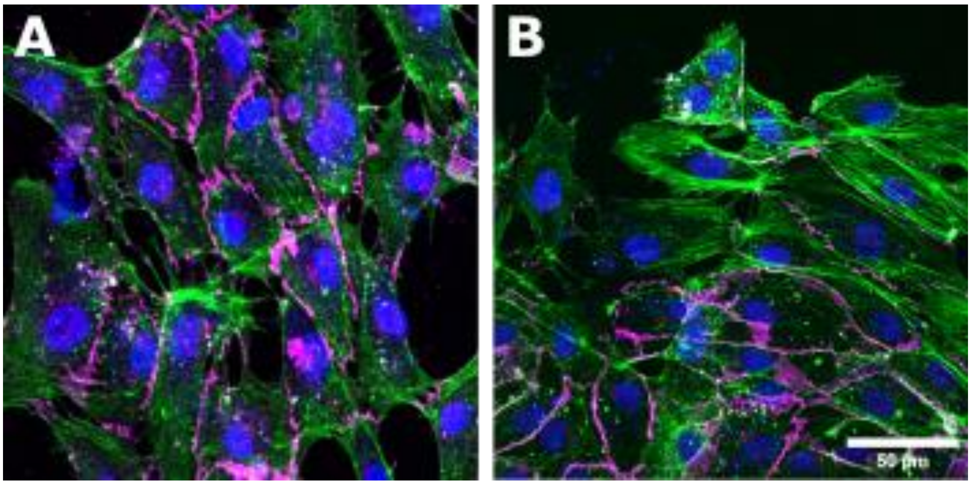


**Figure S3.** Endothelium formation on the luminal side of the bilayered collagen material at day 5 (A) and 14 (B). Green: Actin. Blue: DAPI. Magenta: VE-cadherin

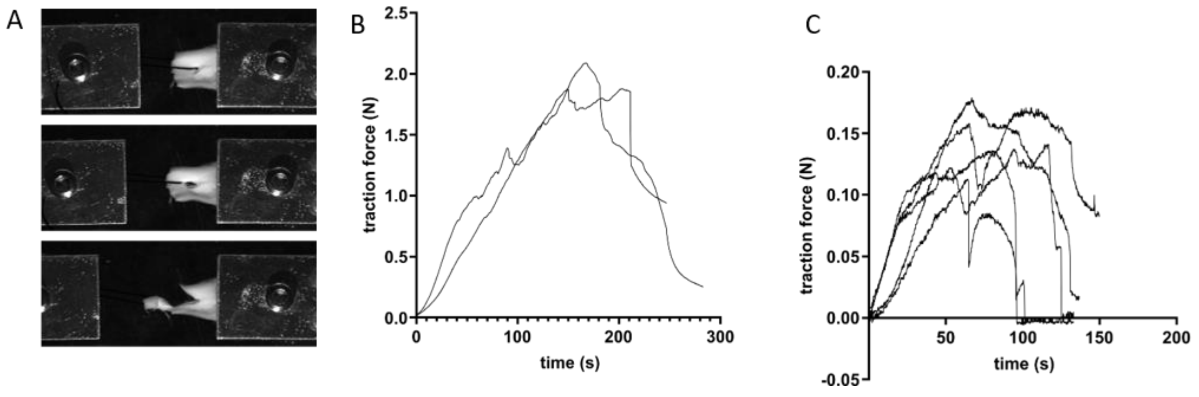


**Figure S4.** Suture retention strength testing. A: Longitudinal strip of the bilayered material fixed at one extremity while the extremity of the suture is displaced to exert pulling. B, C: Measurement of force exerted either on the native artery (B) or the bilayered material (C) with breaking occurring at the maximum force corresponding to the SRS reported value.

**Movie S1.** Inflation test of bilayered collagen tubes with food dye injected into the lumen at internal pressures exceeding 200 mmHg

**Movie S2.** Inflation test of collagen porous material with food dye injected into the lumen**,** leakage through monolayer porous tube is observed below 100 mmHg

**Movie S3.** Uniaxial tensile testing of piglet carotid artery
**Movie S4.** Uniaxial tensile testing of tubular bilayered collagen material
**Movie S5.** Uniaxial tensile testing of tubular non porous collagen material
**Movie S6.** Uniaxial tensile testing of tubular porous collagen material

**Movie S7.** Inflation test of piglet carotid artery under pulsatile pressurization
**Movie S8.** Inflation test of tubular bilayered collagen material under pulsatile pressurization
**Movie S9.** Inflation test of tubular non porous collagen material under pulsatile pressurization
**Movie S10.** Inflation test of tubular porous collagen material under pulsatile pressurization

**Movie S11.** Manipulation of collagen bilayered material with tension exerted on thread sutured at the extremity of the material

**Movie S12.** Suturing test to perform scaffold–bronchus anastomosis, eventually reinforced with TachoSil®
